# Precision-cut liver slices as a model for evaluating therapeutic agents against hepatitis B and delta viruses

**DOI:** 10.64898/2025.12.06.692712

**Authors:** Armando Andres Roca Suarez, Maud Michelet, Marie-Laure Plissonnier, Anaëlle Dubois, Audrey Diederichs, Sarah Heintz, Maria Saez-Palma, Mathieu Ben Abu, Xavier Grand, Anaïs Favre, Alain Chédotal, Nicolaas Van Renne, Simon P. Fletcher, François Mithieux, Pierre Emmanuel Bonnot, Aurélien Dupré, Nicolas Mouton, Michel Rivoire, Sandra Phillips, Elena Palma, Shilpa Chokshi, Fabien Zoulim, Barbara Testoni

## Abstract

**Background:** Developing new therapeutic strategies against hepatitis B virus (HBV) and hepatitis delta virus (HDV) is essential to cure these infections. Yet evaluation of host-targeting agents (HTAs) with *in vitro*/*in vivo* models remains challenging. Therefore, we assessed precision-cut liver slices (PCLS) as an HBV/HDV co-infection model to study virus-host interactions and HTAs.

**Methods:** We generated PCLS from human liver resections and infected them *ex vivo* with HBV and HDV. Tissue viability and architecture were monitored by intracellular ATP, secreted albumin, and multiplexed immunofluorescence. Viral markers were assessed by quantitative PCR, western blot, and light-sheet microscopy in infected slices following treatment with a sodium taurocholate co-transporting polypeptide (NTCP)-targeting peptide, lonafarnib, or the Toll-like receptor 8 agonist selgantolimod (SLGN). Single-cell RNA sequencing was performed to characterize the cellular responses to SLGN.

**Results:** Here we show that our *ex vivo* infection protocol allows the establishment of HBV and HDV infection within the liver’s three-dimensional architecture. Treatment with three well-characterized HTAs produced distinct antiviral effects consistent with each mechanism: an NTCP-targeting peptide blocked entry; lonafarnib induced intracellular hepatitis delta antigen accumulation; SLGN showed anti-HBV activity not previously seen in cultured hepatocytes, indicating cross-talk between immune and infected cells within liver slices.

**Conclusion:** These results provide the characterization of PCLS as an *ex vivo* model of HBV/HDV co-infection and antiviral testing. In the future, PCLS could contribute to expediting pre-clinical characterization of HTAs and to exploring alternatives to animal experimentation.

## INTRODUCTION

Hepatitis B virus (HBV) affects more than 250 million people worldwide and is one of the major etiologies for the development of cirrhosis and hepatocellular carcinoma (HCC).[1] The currently available nucleos(t)ide analogues (NUCs), entecavir and tenofovir, which are direct-acting antivirals (DAAs), require indefinite treatment to maintain viral suppression and prevent the virological relapse occurring when treatment is discontinued.[2] Moreover, HBV co-infection with hepatitis delta virus (HDV) is often associated with a faster liver disease progression in comparison to HBV mono-infection.[3] The entry inhibitor bulevirtide requires prolonged administration or a combination with pegylated interferon alpha (pegIFN-α) to achieve sustained HDV response.[4,5] Thus, HBV and HDV remain unmet medical needs that demand a continuous effort to develop novel treatments and methods for their characterization in order to achieve a viral cure.[6]

The search for new compounds against HBV/HDV is a highly dynamic field that has grown considerably in recent years. This has led to a renewed interest in the development of both novel DAAs and host-targeting agents (HTAs), which opens the possibility to evaluate new combination therapies. However, the pre-clinical evaluation of these new compounds and their combinations is challenging due to the limitations of currently available *in vitro* and *in vivo* study models.[7] Indeed, considering the heterogeneity, proportion, and spatial location of the cell populations composing the liver microenvironment is necessary not only to understand the virus-host interactions in the setting of persistent infection, but equally important for the characterization of potential therapeutic strategies. This is particularly relevant for HTAs directed toward immune cell populations, since, so far, evaluating their antiviral action requires *in vivo* mouse models, which at best imperfectly mirror human innate and adaptive responses,[8] or *in vitro* co-culture systems that lack many of the above-described features found within the human liver.

Aiming to circumvent these limitations and more closely recapitulate the cellular context observed *in vivo*, advanced *ex vivo* models have emerged as a viable alternative to study the liver. In this regard, the precision-cut liver slice (PCLS) model consists of the generation and subsequent *ex vivo* culture of standardized liver sections.[9] PCLS have the advantage of retaining the complex multi-cellular diversity and architecture of the hepatic microenvironment, while offering the practical aspects of an *in vitro* model. In this context, PCLS have been employed as a model for a wide variety of liver pathologies that include hepatitis C virus (HCV) infection, metabolic liver disease, and HCC.[10,11] However, the permissiveness of this model to HBV and HDV infection has not been reported.

Thus, in this study, we evaluate PCLS as an *ex vivo* HBV/HDV infection model, as well as its pertinence for the pre-clinical characterization of HTAs against these viruses.

## METHODS

### Preparation of human PCLS

Human liver tissue for PCLS preparation was obtained in collaboration with the Biological Resources Center and the surgical department of the Centre Léon Bérard (CLB), and the department of surgical oncology of the Hôpital Privé Jean Mermoz. Clinical data were collected by the Ex Vivo Platform of the Cancer Research Center of Lyon (CRCL) (**Table S1**). Liver donors were selected according to the following exclusion criteria: 1) clinical history of HBV or HCV infection, 2) diagnosis of liver fibrosis/cirrhosis or steatosis, and 3) chemotherapy/immunotherapy within three weeks prior to surgical intervention. Collected human liver tissues were kept in sterile transport solution, composed of 1X PBS (CS1PBS00, Eurobio Scientific, Paris, FR), 50 U/mL of penicillin/streptomycin, and 100 µg/mL gentamicin (15140122, 15710049, Gibco, Billings, MT, US). Liver tissue cores (6 mm in diameter) were prepared using biopsy punches (BP-60F, Kai Medical, Honolulu, HI, US) and embedded into 3% ultrapure low melting point agarose blocks (16520050, Thermo Fisher Scientific, Waltham, MA, US). Liver cores were further processed into PCLS (250 µm) using a VT1200 vibratome (Leica Biosystems, Nußloch, DE). Slicing solution was composed of Williams’ E medium (22551-022, Gibco) supplemented with 50 U/mL of penicillin/streptomycin. Slices were incubated individually during a recovery step (37°C, 20 rocks/min, 2 h) in 24-well plates containing PCLS culture medium: Williams’ E medium supplemented with 5% FCII serum (SH30066.02, Cytiva, Marlborough, MA, US), 50 U/mL of penicillin/streptomycin, 15 mM HEPES (15630080, Gibco) and 1X Insulin-Transferrin-Selenium (41400045, Gibco). Subsequently, PCLS were placed on Millicell culture inserts (PICM01250, Merck KGaA, Darmstadt, DE) containing 450 µL of PCLS medium, as previously described.[12] PCLS medium was changed daily and viability evaluated by the quantification of intracellular ATP (CellTiter-Glo 2.0, G9241, Promega, Madison, WI, US) and extracellular albumin (EHALB, Thermo Fisher Scientific).

### *Ex vivo* PCLS infection with HBV/HDV and treatment with HTAs

HBV and HDV viral stocks employed for the *ex vivo* infection of PCLS were prepared as previously described.[13,14] Cell number in each slice was estimated as previously reported,[11] in order to perform infection with 400 and 60 viral genome equivalents (vge)/cell of HBV and HDV, respectively. Briefly, PCLS were placed in 24-well plates without cell culture inserts and infected overnight using PCLS medium containing 4% polyethylene glycol (PEG). Subsequently, PCLS were washed four times with 1X PBS and placed back on cell culture inserts. Alternatively, PCLS were infected with UV-inactivated HBV (3300 µW/cm^2^ for 30 min) using the same method described above. A myristoylated preS1-peptide (i.e., GTNLSVPNPLGFFPDHQLDPAFRANSNNPDWDFNPNKDHWPEANKVG) used as entry inhibitor (EI) was synthesized by GenScript (Rijswijk, NL). PCLS were treated with EI (1 µM) for 24 h prior to HBV/HDV infection and maintained with the compound (100 nM) post-infection. Alternatively, PCLS were treated with selgantolimod (SLGN, 1 µM, 48 h, Gilead Sciences, Foster City, CA, US), lonafarnib (LN, 2 µM, 48 h, SML1457, Sigma-Aldrich, St. Louis, MO, US) or 3TC (20 µM, 144 h, L1295-10MG, Sigma-Aldrich) post-infection. Uninfected PCLS employed for scRNA-seq were treated with SLGN (1 µM, 48 h) or control (dimethyl sulfoxide, DMSO). Medium was changed daily for all conditions.

### Nucleic acid extraction and quantification

PCLS were lysed for the extraction of HBV nucleic acids using the MasterPure DNA and RNA purification kit (MC85200, LGC Biosearch Technologies, Petaluma, CA, US) according to the manufacturer’s instructions. Samples were processed following the ICE-HBV guidelines,[15] using TaqMan Master Mix (4364338, Applied Biosystems, Waltham, MA, US). Covalently closed circular (ccc)DNA amplification was performed on RNase A-and Exo I/III-treated samples in order to degrade HBV RNA, genomic DNA and incomplete double-stranded relaxed circular (rc)DNA intermediate species (MRNA092, X40520K, EX4425K, LGC Biosearch Technologies). Hemoglobin subunit beta (*HBB*) served as internal reference for HBV cccDNA and total DNA quantification; glucuronidase beta (*GUSB*) was used for total RNA. All reactions were carried out with the QuantStudio 7 Flex System (Thermo Fisher Scientific). PCLS were lysed for the extraction of HDV RNA using the Monarch Total RNA Miniprep kit (T2010S, New England Biolabs, Ipswich, MA, US). Total HDV RNA quantification was performed using SYBR Green Master Mix (A25741, Applied Biosystems) and a LightCycler 480 instrument (Roche Diagnostics, Basel, CH). Quantification of genomic and anti-genomic HDV RNA was performed as previously described,[16] employing QX200 droplet digital (dd)PCR EvaGreen Supermix (186-4033, Bio-Rad, Hercules, CA, US). Samples were partitioned into the Automated Droplet Generator with specific oil for probes, amplified in the C1000 Touch thermal cycler, and analyzed with the QX200 Droplet Reader using QuantaSoft software (v1.7.4, Bio-Rad). Primer sequences can be found in **Table S2**.

### Western blotting

PCLS were lysed in RIPA buffer supplemented with protease inhibitor cocktail (PIC, 05056489001, Roche Diagnostics). Protein samples (60 µg) were loaded in 12% polyacrylamide gels (4561043, Bio-Rad) and run at 200 V for 40 min, followed by transfer to nitrocellulose membranes (1704159, Bio-Rad). Membranes were incubated with primary antibodies against β-actin (ab6276, 1/5000, Abcam, Cambridge, UK) and hepatitis delta antigen (HDAg, 1/500), followed by three washes with 1X PBS and incubation with secondary antibodies (1/5000, 12-348, 12-349, Sigma-Aldrich). Blots were revealed using a ChemiDoc MP Imaging System (Bio-Rad).

### Cytokine quantification

Cytokine profiles in the supernatant of PCLS were characterized by IsoPlexis CodePlex secretome assay, using the Innate Immune Panel (S-PANEL-2L03-4) and an IsoSpark instrument (Bruker Cellular Analysis, Billerica, MA, US).

### Immunofluorescence

PCLS (days 1, 3, and 5 post-cutting) were fixed in 10% buffered formalin and embedded in paraffin. Tissue sections (3-µm-thick) were prepared and fully automated multiplexed detection was performed on a Leica Bond RX automated immunostainer (Leica Biosystems). Sequential immunofluorescence for albumin (hepatocytes), CD68 (general myeloid), CD3 (T cells), and CD20 (B cells) was carried out with the OPAL 6-Plex detection kit (NEL821001KT, Akoya Biosciences, Marlborough, MA, US), which employs tyramide signal amplification (TSA)-conjugated fluorophores. Slides were dewaxed at 72°C with Bond Dewax Solution (AR9222, Leica Biosystems) and antigens retrieved using Bond Epitope Retrieval Solution 1 (pH 6, AR9961, Leica Biosystems) for 20 min at 97°C. In a first step for CD20 staining, slides were incubated with mouse monoclonal anti-CD20 for 30 min (M0755, 1/600, Agilent, Santa Clara, CA, US), and incubated with rabbit monoclonal anti-mouse secondary antibody for 30 min (OPAL 520, 1/150, Leica Biosystems). Slides were then sequentially incubated with anti-albumin (A0001, 1/20000, Agilent) for 30 min and anti-rabbit secondary antibody OPAL 690 (1/150); anti-CD68 (IR61361-2, 1/50, Agilent) and anti-mouse secondary antibody OPAL 570 (1/150) for 30 min; anti-CD3 (IR503, 1/100, Agilent) for 60 min and anti-rabbit secondary antibody OPAL 480 (1/150) for 30 min. Slides were counterstained with DAPI (nuclear DNA) and then mounted using ProLong Gold Antifade Reagent (P36930, Thermo Fisher Scientific). All sections were scanned using a PhenoImager HT instrument (Akoya Biosciences) at a 20× magnification factor. The whole slide views for the native images were generated using Phenochart (v1.1, Akoya Biosciences).

### 3D imaging by light-sheet microscopy

#### Bleaching

To remove pigmentation and reduce background, tissue bleaching was carried out as previously described.[17] PCLS were dehydrated for 1 h at room temperature (RT) in ascending concentrations of methanol (MeOH, 20847.320, VWR, Radnor, PA, US) in 1X PBS (20, 40, 60, and 80%). PCLS were placed overnight under light and rolling agitation with a 6% hydrogen peroxide solution (216763-500ML, Sigma-Aldrich) in 100% MeOH. The next day, PCLS were rehydrated for 1 h at RT in descending concentrations of MeOH (80, 60, 40, and 20%), washed three times with 1X PBS, and blocked in 1X PBS-0.2% Gelatin (24350.262, VWR)-0.5% Triton X-100 (X100-1L, Sigma-Aldrich) (PBSGT0.5%) solution for 3 days.

#### Whole-mount immunolabeling

PCLS were transferred to a solution containing the primary antibodies diluted in PBSGT0.5% and incubated at 37°C with agitation at 20 rpm for 7 days (hepatitis B virus surface antigen, HBsAg: BSB5622, 1/200, BioSB, Santa Barbara, CA, US; HDAg: EDHD001, 1/500, Kerafast, Boston, MA, US). This was followed by six washes of 1 h in PBSGT0.5% at RT. Then, secondary antibodies were diluted in PBSGT0.5% and passed through a 0.22-µm filter. PCLS were incubated at 37°C with agitation at 20 rpm in the secondary antibody solution for 7 days (anti-Mouse IgG H&L-AF555: ab150110, 1/800, Abcam, Cambridge, UK; Helix NP Green (nuclear DNA): 425303, 1/1000, BioLegend, San Diego, CA, US) and washed six times for 1 h in PBSGT0.5% at RT.

#### Agarose embedding

To facilitate the handling and imaging of immune-labeled samples with the light-sheet microscope, PCLS were embedded prior to clearing in 1.5% agarose (2267.5, Carl Roth GmbH + Co, Karlsruhe, DE). Agarose was dissolved in 1X TAE and heated in a microwave oven until a homogeneous solution was obtained. PCLS were placed in polystyrene weighing dishes and cooled agarose was poured over them, taking care to avoid introducing air bubbles. After removing the PCLS embedded in agarose from the mold, agarose was trimmed as close as possible to the sample with a razor blade.

#### Sample clearing

The iDISCO+ protocol was performed.[18] After immunolabeling and embedding, samples were placed in TPP tubes, dehydrated for 90 min in ascending concentrations of MeOH in 1X PBS (20, 40, 60, 80, and 100% twice) under rotating agitation. Samples were next incubated in a solution of 67% dichloromethane (DCM, 5895812000, Sigma-Aldrich) and 33% MeOH overnight, followed by 100% DCM for 45 min at RT under rotating agitation. Finally, samples were put in 100% dibenzyl ether (DBE, 108014-1kg, Sigma-Aldrich) for a minimum of 2 days.

#### 3D imaging

Cleared samples were imaged at the 3D HUMAN facility, with a Blaze light-sheet microscope (Miltenyi Biotec, Bergisch Gladbach, DE) equipped with an sCMOS camera 4.2MP (2048 × 2048) controlled by the ImSpector Pro acquisition software (v8.0.2, Miltenyi Biotec). The light-sheet, of 4-µm thickness, was generated by lasers at three different wavelengths (i.e., 488, 561, and 639 nm). A 4× objective with 1.66× magnification lenses was used. Samples were supported by a sample holder (Miltenyi Biotec), placed in a tank filled with DBE, and illuminated by the laser light-sheet from one side. Fast Tiling Scan and LightSpeed Mode were used, respectively, to shift the sample inside the light-sheet while imaging and speed up image acquisition to obtain sharper images. The images were acquired in 16-bit TIFF format. Mosaics of image tiles were assembled with an overlap of 20% between the tiles by MACS iQ View image processing software (v1.2.4, Miltenyi Biotec), and then converted to Imaris 3D images.

### Single-cell RNA sequencing

Following *ex vivo* culture, PCLS were snap-frozen in liquid nitrogen and stored at −80°C until tissue processing. In order to obtain 25 mg of tissue, five liver slices per condition were pooled, fixed with 4% paraformaldehyde for 20 h (BP531-25, Thermo Fisher Scientific), dissociated using 1 mg/mL Liberase TH for 20 min (5401135001, Merck KGaA) followed by mechanical dissociation with spring scissors, and preserved according to the Tissue Fixation & Dissociation for Chromium Fixed RNA Profiling protocol CG000553 (10x Genomics, Pleasanton, CA, US).

Cell number was determined to obtain an expected recovery of 10000 cells/channel, and samples were run on a Chromium Controller system (10x Genomics) according to the manufacturer’s instructions. Single-cell (sc)RNA-seq libraries were generated with the Chromium Fixed RNA Profiling Kit for Multiplexed Samples (10x Genomics, Human Transcriptome Probe Set v1.0.1), and sequenced on a NovaSeq 6000 platform (Illumina, San Diego, CA, US) to obtain 50000 reads/cell.

Data analysis was performed as described in Van Renne *et al*., JHEP Rep. 2026.[19] Briefly, matrices were loaded into Seurat (v5.0.2)[20] using the *Read10X* function. Cells were filtered based on mitochondrial content (<25%), total number of transcripts (>1500, <99999), and total number of genes (>200). Data were normalized using the *NormalizeData* function to logtransform the counts. Variable features were identified with the *FindVariableFeatures* function, selecting the top 2000 most variable genes. Data were then scaled using the *ScaleData* function. Principal component analysis (PCA) was performed using *RunPCA* on the first 30 dimensions. Harmony integration was performed to correct for donor effects.[21] The *RunUMAP* function was used for dimensionality reduction using the first 30 dimensions. To identify cell clusters, the *FindNeighbors* function was employed, followed by *FindClusters* with a resolution of 3. Clusters were annotated using canonical cell type markers.[22] Gene signature scores were estimated using fast gene set enrichment analysis (fgsea, v1.39.4),[23] Hallmark, Reactome, and Gene Ontology gene sets from the MSigDB (v2026.1),[24], as well as the SLGN-UP and SLGN-DOWN signatures described in Roca Suarez, Plissonnier *et al*., Gut 2024.[25] Graphs were generated using the *DimPlot, FeaturePlot, VlnPlot*, and *DotPlot* functions of Seurat. ScRNA-seq data from healthy liver tissues and code for the re-analyses presented in **Fig. 2B** and **Fig. S3** were obtained from Guilliams *et al*., Cell 2022 (livercellatlas.org).[22]

### Statistics and reproducibility

All statistical analyses were performed in Prism (v10, Dotmatics, Boston, MA, US). Tests used are indicated in the figure legends (Kruskal-Wallis with Dunn’s multiple comparison test, analysis of variance with Dunnett’s multiple comparison test, and Mann-Whitney test). Statistical significance was defined as *p* < 0.05 or FDR < 0.05. *n* numbers denote individual liver tissue donors (colorcode). Biological replicates are defined as individual liver slices from the same donor and experimental condition.

## RESULTS

### PCLS are permissive to *ex vivo* HBV and HDV infection

In order to assess the potential use of PCLS as an *ex vivo* HBV/HDV infection model, we established an experimental protocol consisting of PCLS preparation and their subsequent mono- or co-infection with HBV and HDV for a five-day period (**Fig. 1A**). Clinical data from the liver tissue donors can be found in **Table S1**. PCLS viability was monitored daily by the quantification of intracellular ATP and extracellular albumin (**Fig. 1B, C**). In this experimental setting we were able to detect total HBV RNA and cccDNA in PCLS infected with HBV and co-infected with HDV (**Fig. 1D, E**), as well as total HDV RNA and HDAg in PCLS mono- or co-infected with HDV (**Fig. 1F, G**). Moreover, immunostaining combined with tissue bleaching and light-sheet microscopy allowed detection of a positive signal for HBsAg and HDAg in HBV/HDV-infected PCLS (**Fig. S1** and **Fig. S2**). Active viral replication was confirmed by the quantification of total HBV RNA and cccDNA levels in comparison to a UV-inactivated HBV inoculum (**Fig. 1H, I**) and by the detection of both genomic and anti-genomic HDV RNA species by specific ddPCR (**Fig. 1J**). These results suggest that viral replication is indeed taking place following *ex vivo* infection, and that PCLS are thus permissive to both HBV and HDV.

**Fig 1.**
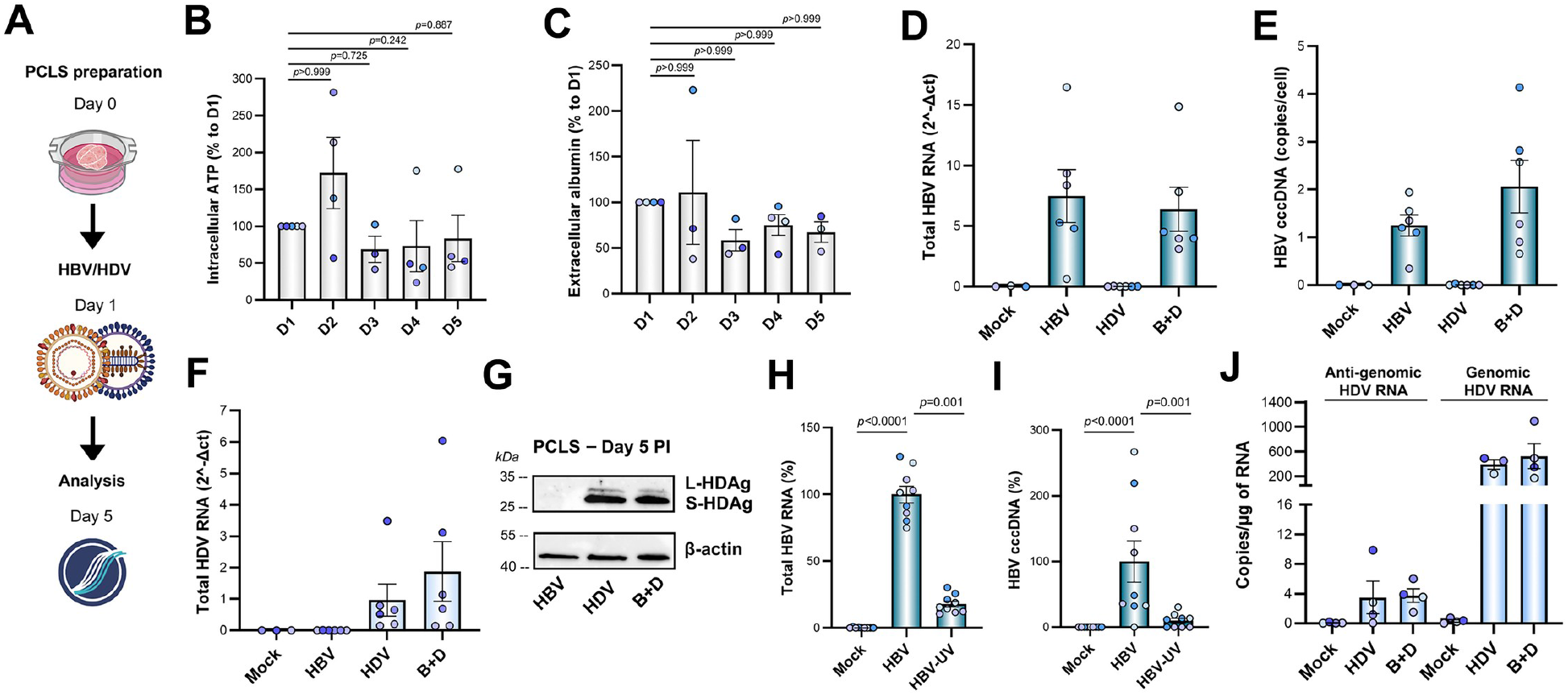
PCLS are permissive to *ex vivo* HBV and HDV infection. **(A)** Experimental protocol consisting of the preparation of PCLS, their mono- or co-infection with HBV/HDV for a five-day period and the quantification of viral parameters. Image created in BioRender. **(B–C)** Quantification of (B) intracellular ATP and (C) extracellular albumin in PCLS maintained in culture for five days (*n* = 4, Kruskal-Wallis test). **(D–F)** Quantification of (D) total HBV RNA, (E) cccDNA and (F) total HDV RNA by qPCR in HBV/HDV mono- or co-infected PCLS (*n* = 3). **(G)** Detection by western blot of intracellular large and small HDAg in HDV-or HBV/HDV-infected PCLS. **(H–I)** Quantification of (H) total HBV RNA and (I) cccDNA by qPCR in PCLS infected with HBV or UV-inactivated HBV (*n* = 3, Kruskal-Wallis test). **(J)** Quantification of genomic and anti-genomic HDV RNA by ddPCR (*n* = 3). Bars represent mean ± SEM. Dot colors indicate individual liver tissue donors (*n*). cccDNA, covalently closed circular DNA; ddPCR, droplet digital PCR; HBV, hepatitis B virus; HDAg, hepatitis delta antigen; HDV, hepatitis delta virus; PCLS, precision-cut liver slices.

**Fig 2.**
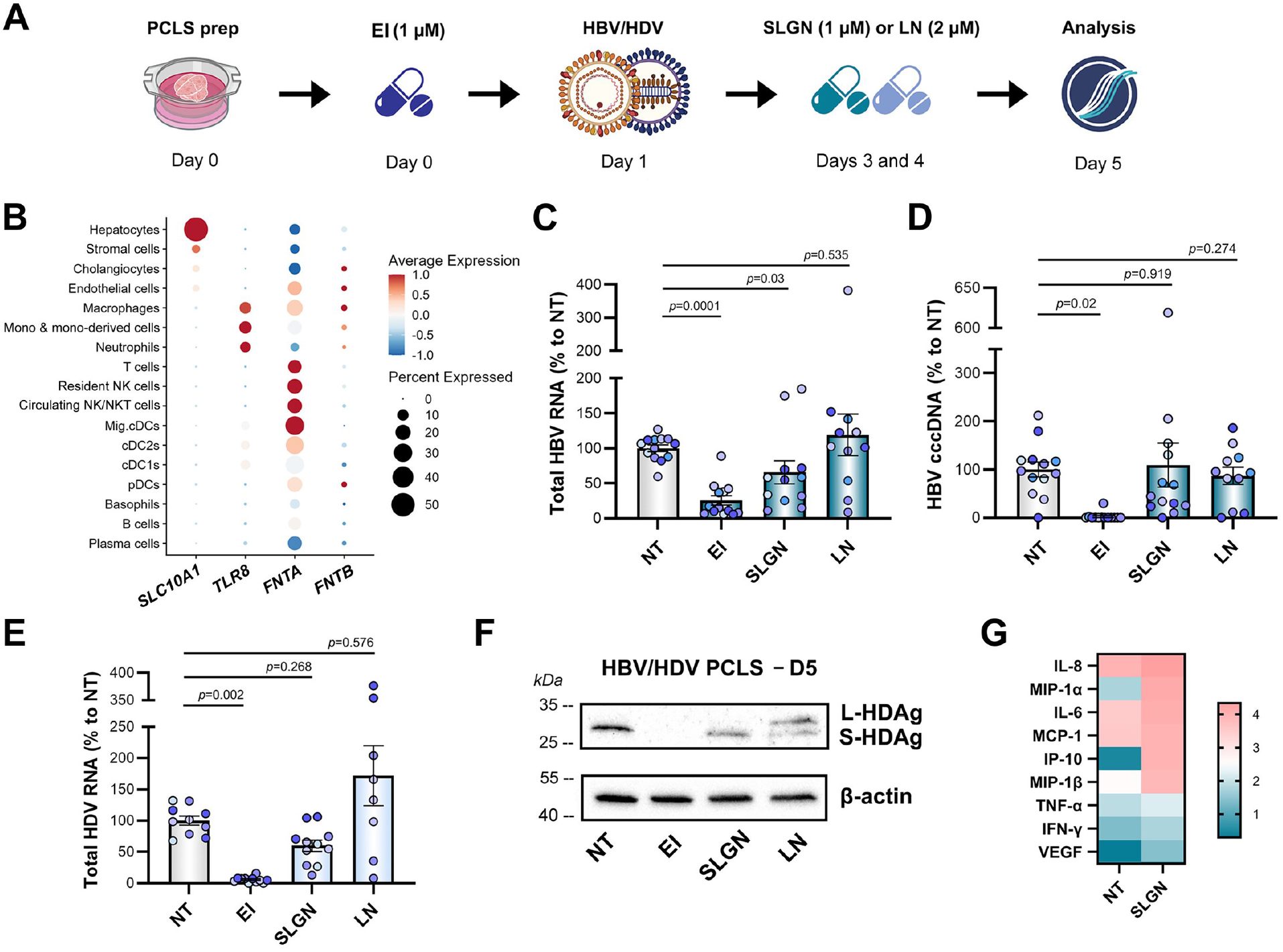
PCLS allow the evaluation of HTAs against HBV/HDV infection. **(A)** Experimental protocol consisting of the preparation of PCLS, the use of EI (1 µM) prior to HBV/HDV infection, and LN (2 µM) or SLGN (1 µM) post-infection, in order to evaluate their antiviral effect by the quantification of viral parameters. Image created in BioRender. **(B)** Cell populations in the human liver microenvironment showing expression of *SLC10A1* (NTCP, EI target), *TLR8* (SLGN target), *FNTA* and *FNTB* (LN targets). ScRNA-seq data obtained from Guilliams *et al*., Cell 2022.[22] **(C)** Quantification of total HBV RNA by qPCR in HBV/HDV-infected PCLS treated with HTAs (*n* = 5, one-way ANOVA). **(D)** Quantification of HBV cccDNA by qPCR in HBV/HDV-infected PCLS treated with HTAs (*n* = 5, one-way ANOVA). **(E)** Quantification of total HDV RNA by qPCR in HBV/HDV-infected PCLS treated with HTAs (*n* = 5, one-way ANOVA). Bars represent mean ± SEM. Dot colors indicate individual liver tissue donors (*n*). **(F)** Detection by western blot of intracellular large and small HDAg in HBV/HDV-infected PCLS treated with HTAs. **(G)** Cytokine levels (log pg/mL) in the supernatant of PCLS treated with SLGN (1 μM, 24 h, *n* = 4), as assessed by IsoPlexis assay. cccDNA, covalently closed circular DNA; cDCs, classical dendritic cells; EI, entry inhibitor; FNTA, farnesyltransferase, CAAX box, subunit alpha; FNTB, farnesyltransferase, CAAX box, subunit beta; HBV, hepatitis B virus; HDAg, hepatitis delta antigen; HDV, hepatitis delta virus; HTAs, host-targeting agents; IFN, interferon; IL, interleukin; IP-10, 10 kDa interferon gamma-induced protein; LN, lonafarnib; MCP-1, monocyte chemoattractant protein 1; Mig.cDCs, migratory cDCs; MIP-1, macrophage inflammatory protein 1; NK cells, natural killer cells; PCLS, precision-cut liver slices; pDCs, plasmacytoid DCs; scRNA-seq, single-cell RNA sequencing; SLC10A1, solute carrier family 10 member 1; SLGN, selgantolimod; TLR8, Toll-like receptor 8; TNF, tumor necrosis factor; VEGF, vascular endothelial growth factor.

### PCLS allow the evaluation of HTAs against HBV/HDV infection

Having determined that PCLS are permissive to HBV and HDV infections, we next explored the utility of this model for evaluating HTAs against these viruses. Using a similar protocol consisting of the analysis of viral parameters five days after inoculation, we additionally included: 1) preinoculation treatment with an EI, which blocks the viral receptor sodium taurocholate cotransporting polypeptide (NTCP) and impairs entry of both viruses, 2) post-inoculation treatment with the selective Toll-like receptor 8 (TLR8) agonist SLGN to induce antiviral responses against HBV through immune cell activation,[25,26] and 3) post-inoculation treatment with the farnesyltransferase inhibitor LN, which disrupts HDV assembly (**Fig. 2A**). Expression levels of the drugs’ cellular targets and their distribution in each population of the liver microenvironment are depicted in **Fig. 2B** and **Fig. S3**, as obtained by the analysis of publicly available scRNA-seq data from human donors.[22] These experiments showed that treatment with EI was able to impair entry of both viruses, as observed by the significant decrease of HBV RNA (*p* = 0.0001) and cccDNA (*p* = 0.02) levels (**Fig. 2C, D**), as well as of HDV RNA (*p* = 0.002) and L-HDAg (**Fig. 2E, F**). SLGN was able to decrease total HBV RNA levels (*p* = 0.03) (**Fig. 2C**), which likely stems from the indirect effect of inflammatory cytokines produced by the immune cell populations present within PCLS (**Fig. 2G** and **Fig. S4**). Moreover, SLGN had no significant impact on cccDNA levels (*p* = 0.919) (**Fig. 2D**), an observation that is in line with previous reports describing the postinfection use of this TLR8 agonist *in vitro*.[26] LN treatment led to the intracellular accumulation of large (L)-HDAg in PCLS (**Fig. 2F**). Finally, considering that NUCs are employed as the backbone of combination therapies evaluating novel HTAs, we assessed their antiviral effect in HBV/HDV-infected PCLS (**Fig. S5A–C**). Our results show that, as expected, treatment with 3TC (lamivudine) decreased total HBV DNA (*p* = 0.002), without affecting the levels of cccDNA (*p* = 0.125) and HDAg (**Fig. S5D–F**). These results suggest that PCLS are a relevant model for the evaluation of antiviral therapies against HBV/HDV infection.

### PCLS as a model to characterize the direct and indirect modulation of the liver microenvironment by HTAs

Aiming to explore the potential usefulness of the model to characterize how HTAs directly and indirectly modulate the cell types present in the human liver microenvironment, we treated PCLS with SLGN (1 µM, 48 h, *n* = 3) and performed scRNA-seq (**Fig. 3A**). This allowed us to observe the heterogeneity of cell populations present within human PCLS, including hepatocytes, cholangiocytes, endothelial cells, fibroblasts/hepatic stellate cells (HSCs), natural killer (NK) cells, T cells, B cells, classical dendritic cells (cDCs), and Kupffer cells (KCs) among others (**Fig. 3B, Fig. S6**, and **Fig. S7**). Considering that KCs and cDCs are the main cell types expressing *TLR8* (**Fig. 3C** and **Fig. S8**), and that KCs are more abundant than cDCs (**Fig. S9**), we focused on KCs to characterize the direct effect of SLGN. Therefore, we employed two gene signatures we had previously identified in response to this TLR8 agonist *in vitro* with human KCs and *in vivo* with uninfected macaques: SLGN-UP, which includes upregulated genes associated with the inflammatory response, and SLGN-DOWN, which includes downregulated genes associated with KC differentiation status.[25] The results of this analysis showed that in line with our previous observations, KCs within uninfected PCLS presented a significant increase of the SLGN-UP signature, in parallel to a decrease of the SLGN-DOWN signature and IFN-α signaling in response to *ex vivo* SLGN treatment (FDR < 0.05) (**Fig. 3D**). Moreover, this is consistent with the selective activation of TLR8 by SLGN, which is mainly mediated by the nuclear factor kappa B (NF-κB) pathway, and the minimal IFN-α production reported in mononuclear phagocytes.[27] This modulation was also observed at the individual gene level by the increase of inflammatory cytokines and the decrease of KC differentiation markers (**Fig. S10**). Moreover, consistent with our previous *in vitro* data showing that IL-6 produced by KCs following SLGN treatment leads to an indirect activation of the inflammatory response and a decrease of metabolic signaling in primary human hepatocytes,[25] we observed the enhanced expression of *IL6* in KCs (FDR < 0.0001) (**Fig. 3E**), a positive enrichment of the IL-6 response pathway in hepatocytes within PCLS, and the downregulation of bile acid metabolism signaling (FDR < 0.05) (**Fig. 3F**). Finally, this indirect modulation of the liver microenvironment could also be appreciated in immune populations that do not express *TLR8*, as NK cells within PCLS showed the activation of multiple signaling pathways associated with the inflammatory response (FDR < 0.05) (**Fig. 3G**). Although not significant, we observed a tendency towards an increased expression of granzyme B (*GZMB*, log2FC = 1.09) and tumor necrosis factor (ligand) superfamily 10 (*TNFSF10*, TRAIL, log2FC = 0.35) following *ex vivo* SLGN treatment (**Fig. S11**). This is consistent with previous reports describing the upregulation of these inflammatory markers in NK cells from peripheral blood mononuclear cells (PBMCs) of healthy individuals or patients with chronic HBV infection treated with SLGN *in vitro*,[27] as well as in NK cells isolated from healthy volunteers following *in vivo* SLGN administration.[28] Overall, these results highlight the potential of PCLS combined with scRNA-seq for characterizing the direct and indirect modulation of the liver microenvironment in response to HTAs.

**Fig 3.**
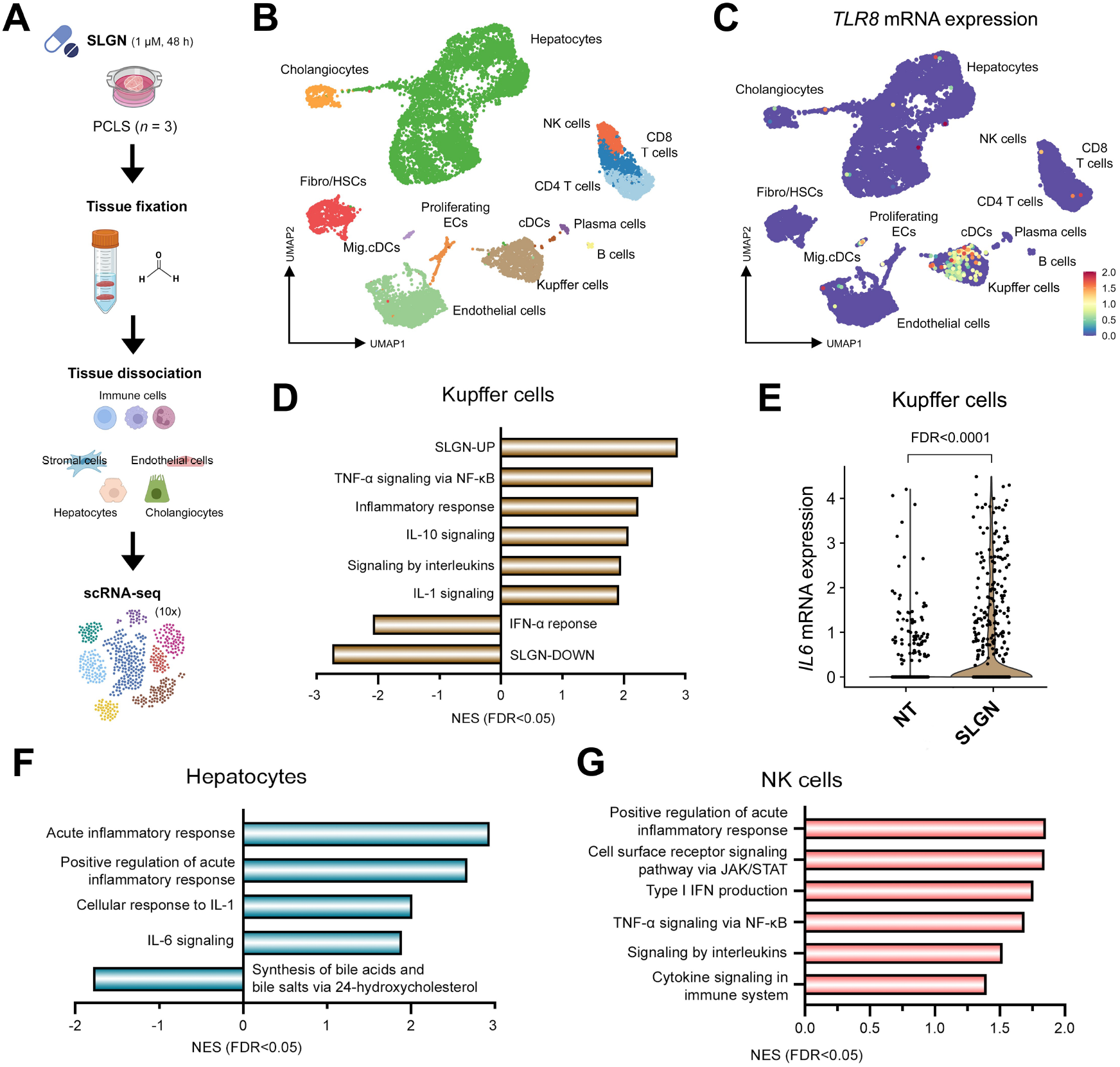
PCLS as a model to characterize the direct and indirect modulation of the liver microenvironment by HTAs. **(A)** Experimental protocol consisting of the preparation of PCLS, the use of SLGN (1 µM, 48 h), tissue fixation, and dissociation in order to perform scRNA-seq (*n* = 3). Image created in BioRender. **(B)** UMAP showing the cell populations identified in PCLS. **(C)** *TLR8* mRNA expression levels in each cell type present within PCLS. **(D)** GSEA showing significantly modulated gene signatures in KCs from PCLS treated with SLGN (FDR < 0.05). **(E)** *IL6* mRNA expression levels in KCs from PCLS treated with SLGN (FDR < 0.0001). **(F)** GSEA showing significantly modulated gene signatures in hepatocytes from PCLS treated with SLGN (FDR < 0.05). **(G)** GSEA showing significantly modulated gene signatures in NK cells from PCLS treated with SLGN (FDR < 0.05). *n* numbers denote individual liver tissue donors. cDCs, classical dendritic cells; ECs, endothelial cells; GSEA, gene set enrichment analysis; HSCs, hepatic stellate cells; HTAs, host-targeting agents; IFN, interferon; IL, interleukin; JAK, Janus kinase; KCs, Kupffer cells; Mig.cDCs, migratory cDCs; NES, normalized enrichment score; NF-κB, nuclear factor kappa B; NK cells, natural killer cells; PCLS, precision-cut liver slices; scRNA-seq, singlecell RNA sequencing; SLGN, selgantolimod; STAT, signal transducer and activator of transcription; TLR8, Toll-like receptor 8; TNF, tumor necrosis factor; UMAP, uniform manifold approximation and projection.

## DISCUSSION

HBV/HDV infection and its associated complications still represent a considerable health burden worldwide.[1] This, together with the limitations of current pre-clinical models for the characterization of potential therapeutic strategies, is driving the field to explore new directions. In this context, our results have several implications with respect to the use of PCLS as an *ex vivo* study model for virally induced liver disease.

Firstly, we report that PCLS are permissive to HBV and HDV infection (**Fig. 1**). Our observations highlight the applicability of this model to studying the basic biology of hepatotropic viruses in a cellular context closer to the human liver microenvironment. This provides a significant added value compared to studies performed with isolated cells, conditioned media, co-cultures, and the currently available animal models, including liver-humanized mice.[7]

Secondly, PCLS reproduced the expected pharmacology of mechanistically distinct HTAs (**Fig. 2**). This suggests that PCLS could represent a valuable tool to complement the use of *in vitro*/*in vivo* models, such as woodchuck hepatitis virus (WHV)-infected animals,[29] by providing antiviral profiles in order to further strengthen the rationale to move forward with the clinical evaluation of a given compound.

Thirdly, the presence of both parenchymal and non-parenchymal cells within PCLS makes it possible to mimic the interactions these hepatic populations could have in response to immunomodulatory agents (**Fig. 3**). This can be observed, for instance, with SLGN administration, as this compound has been reported not to have a direct antiviral effect on HBV-infected hepatocytes, and instead to decrease viral parameters via the activation of immune cell populations.[25–27] Moreover, the increased levels of IL-6, IL-8 (*CXCL8*), MCP-1 (*CCL2*), MIP-1β (*CCL4*), IP-10 (*CXCL10*), and IFN-γ are consistent with those observed during the clinical evaluation of this TLR8 agonist.[30–32] Several of these soluble factors, including IL-6, IFN-γ, and TNF-α, have been reported to induce a decrease of viral markers in infected hepatocytes.[25,33,34] Thus, the PCLS model offers the possibility to investigate cellular crosstalk within the liver microenvironment.

Finally, from an ethical point of view, there has been a recent push to accelerate the development of new approach methodologies (NAMs) in order to replace, reduce and refine (3Rs) the use of animals.[35,36] Human-based models such as PCLS have the potential to offer an alternative for the study of disease pathobiology and potential therapeutic compounds, while addressing these ethical concerns.

Nonetheless, further technical improvements of the model would need to be implemented in order to prolong the viability of PCLS beyond the infection period employed in our work. This would open the possibility to study questions that require longer infection conditions, such as the interference of HDV during the HBV cycle.[37] Additional limitations are the dependence on the availability of freshly resected human specimens and the inter-experimental variability linked to donor heterogeneity.[38] Equally important would be to determine the impact of a donor’s clinical history on the susceptibility of PCLS to *ex vivo* HBV/HDV infection and therapeutic interventions, as well as how this reflects the situation taking place in the healthy human liver.

In conclusion, we show that PCLS are permissive to *ex vivo* HBV/HDV infection and respond to HTAs in a manner consistent with their known mechanisms. PCLS therefore have the potential to expedite the pre-clinical characterization of HTAs and to contribute to the development of alternatives to animal experimentation.

## Contributors

Conceptualization: FZ, BT. Methodology: AARS, MM, MLP, ADu, ADi, SH, MSP, MBA, XG, AF, AC, NVR. Investigation: AARS, MM, MLP, ADu, ADi, SH, MSP, MBA, XG, AF, AC, NVR. Visualization: AARS, MM. Supervision: FZ, BT. Resources: SPF, FM, PEB, ADup, NM, MR, SP, EP, SC. Funding acquisition: FZ, BT. Writing – original draft: AARS. Writing – review & editing: All authors.

## Patient and public involvement

Patients and/or the public were not involved in the design, or conduct, or reporting, or dissemination plans of this research.

## Data availability statement

Data related to the detection of viral antigens by light-sheet microscopy (**Fig. S1** and **Fig. S2**), hepatic expression and distribution of cellular targets with a relevance to this work (**Fig. S3**), the characterization of PCLS by immunofluorescence (**Fig. S4**), the effect of 3TC over viral parameters (**Fig. S5**), scRNA-seq analyses (**Fig. S6–11**), clinical data of each liver tissue donor (**Table S1**), and primer sequences (**Table S2**) can be found in the *Supplementary Information*. Raw count matrices, a processed Seurat object, and code to reproduce the analyses related to the characterization of SLGN-or control (DMSO)-treated PCLS (*n* = 3) by scRNA-seq are publicly available to download at Zenodo (https://doi.org/10.5281/zenodo.22143679).

## Ethics statements

Patient consent for publication: Not applicable.

Ethics approval: Human liver tissue for PCLS preparation was obtained in collaboration with the Biological Resources Center and the surgical department of the CLB, and the department of surgical oncology of the Hôpital Privé Jean Mermoz (French ministerial authorizations BB-0033-00050, AC-2008-238, and DC-2025-7445). Non-opposition for the use of the samples was obtained from all patients.

## Acknowledgments

The authors would like to thank Pélagie Huchon, Claire Verzeroli, Charlotte Hernandez, and Jennifer Molle for the preparation of PCLS; Elisabeth Laurendon, Sophie Léon, and Séverine Tabone-Eglinger from the Ex Vivo Platform of the CRCL for the anonymization of clinical data; Cyril Dégletagne from the Cancer Genomics Platform of the CRCL for technical advice and conducting scRNA-seq experiments; Dr. Birke Bartosch for providing the anti-HDAg antibody; Dr. Julien Courchet of the Institut NeuroMyoGène for technical advice.

