## Supplementary information for "Precision-cut liver slices as a model for evaluating therapeutic agents against hepatitis B and delta viruses"

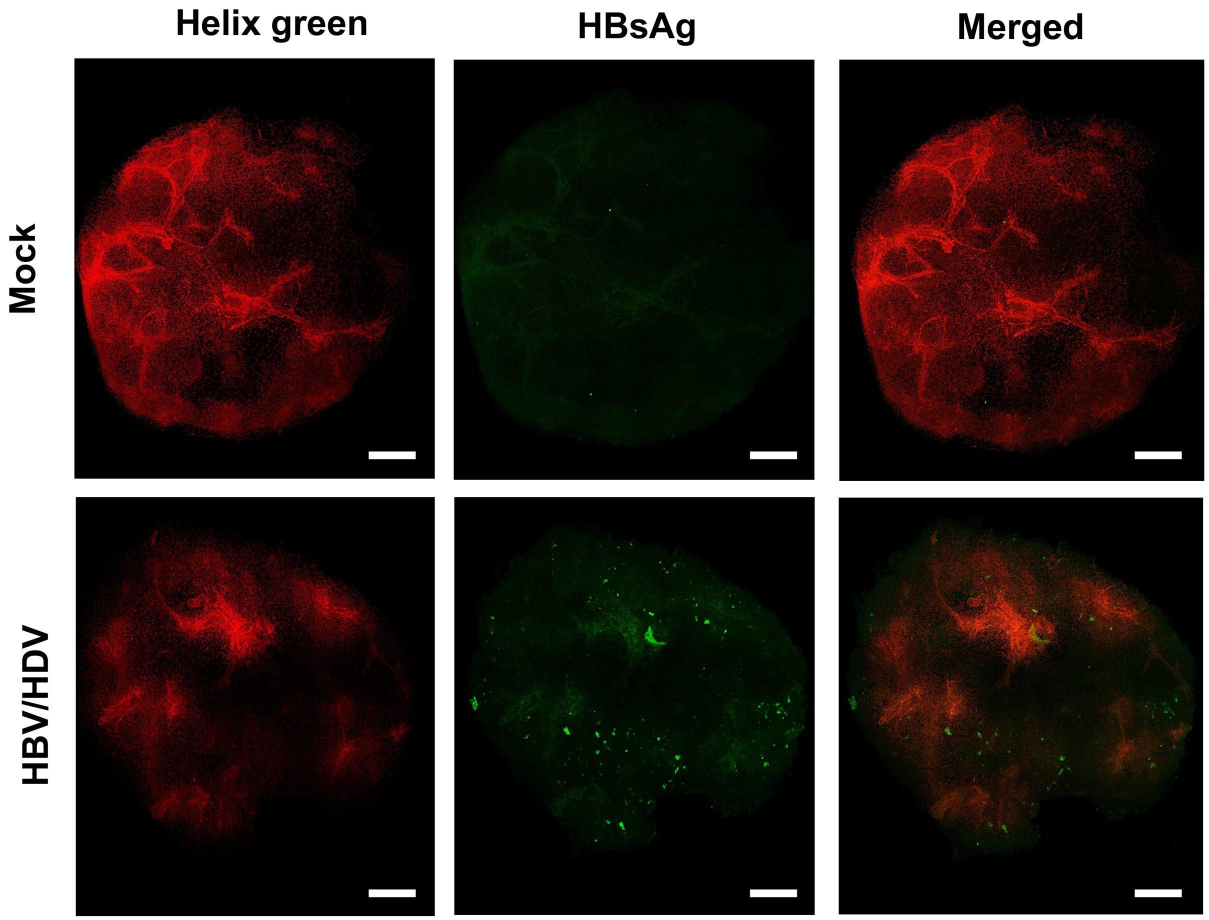

**Fig. S1. Detection of HBsAg in HBV/HDV-infected PCLS.** Combination of immunostaining, tissue bleaching, and light-sheet microscopy for the detection of viral antigens in PCLS at day 5 post-infection. Scale bar represents 500 µm. Helix green = nuclear DNA. HBsAg, hepatitis B virus surface antigen; HBV, hepatitis B virus; HDV, hepatitis delta virus; PCLS, precision-cut liver slices.

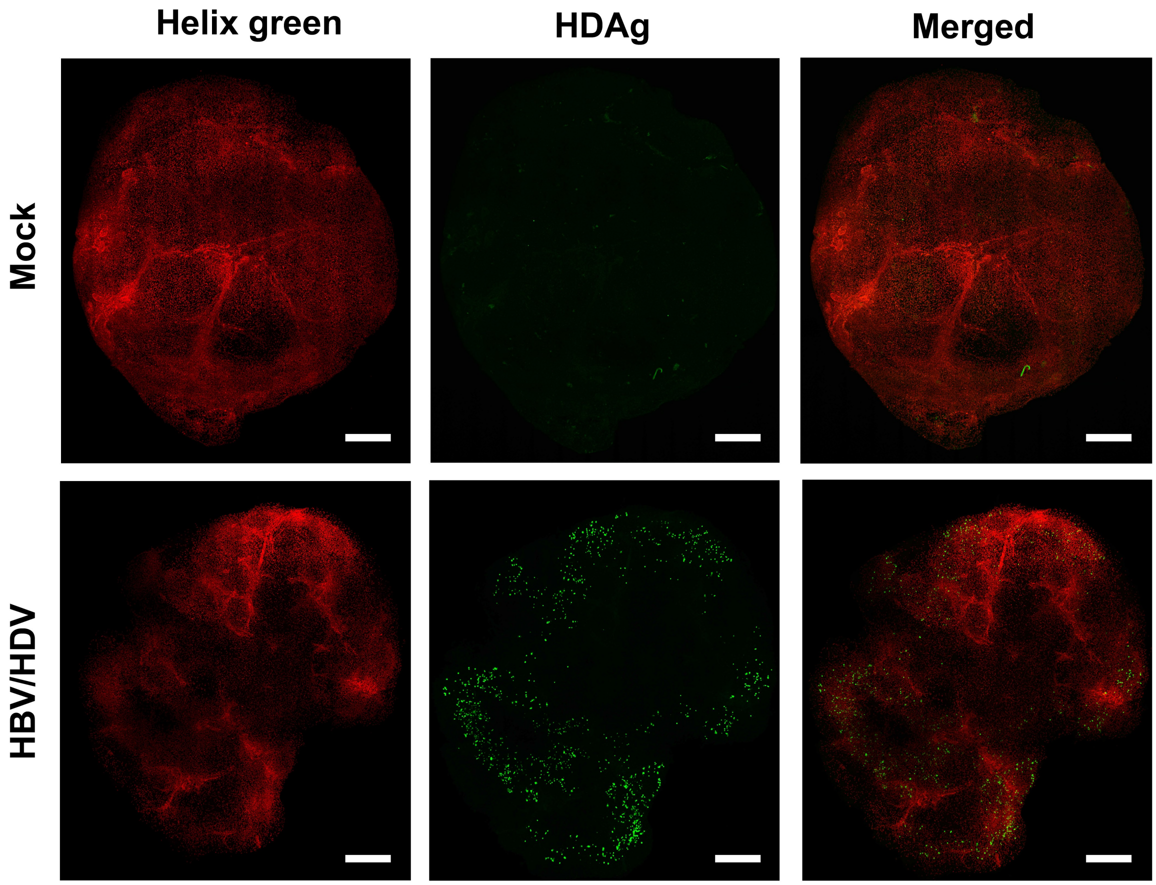

**Fig. S2. Detection of HDAg in HBV/HDV-infected PCLS.** Combination of immunostaining, tissue bleaching, and light-sheet microscopy for the detection of viral antigens in PCLS at day 5 post-infection. Scale bar represents 500 µm. Helix green = nuclear DNA. HBV, hepatitis B virus; HDAg, hepatitis delta antigen; HDV, hepatitis delta virus; PCLS, precision-cut liver slices.

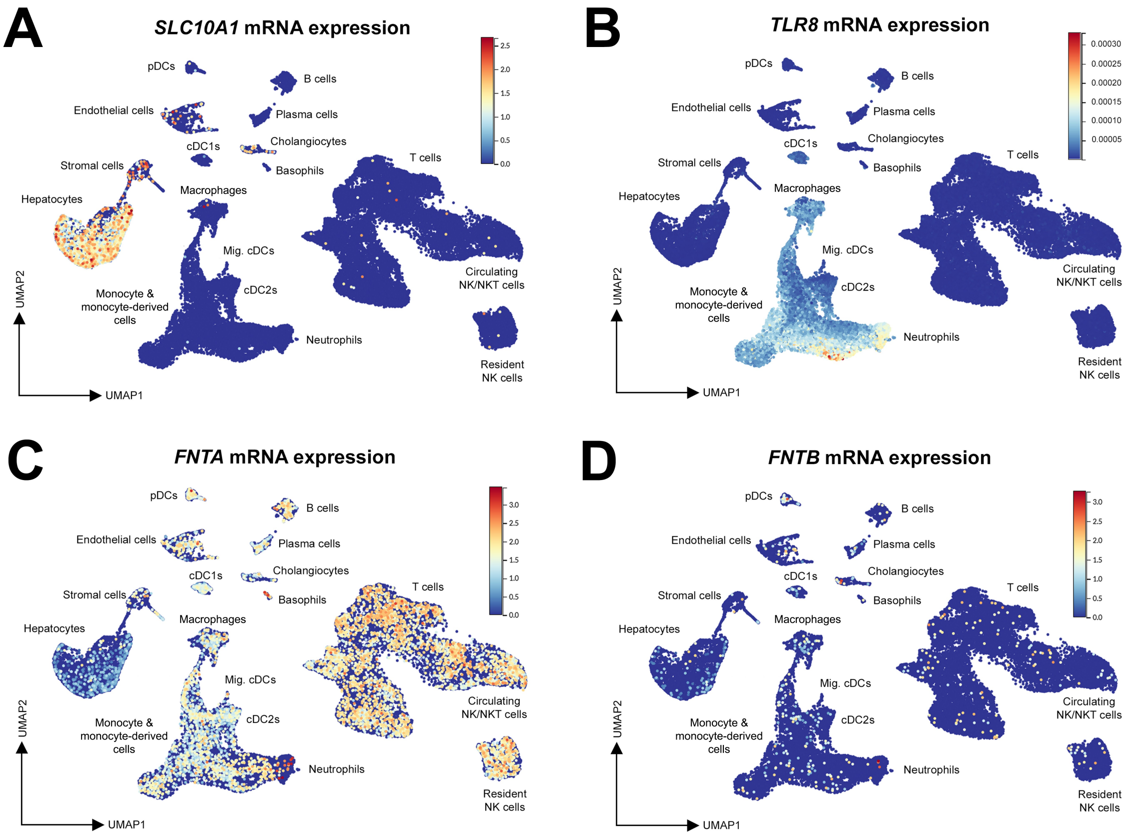

**Fig. S3. Hepatic expression of cellular components of relevance to the HTAs employed. (A–D)** Cell populations in the human liver microenvironment showing expression of (A) *SLC10A1* (NTCP, EI target), (B) *TLR8* (SLGN target), (C–D) *FNTA* and *FNTB* (LN targets). ScRNA-seq data obtained from Guilliams *et al.*, Cell 2022.[1] cDCs, classical dendritic cells; EI, entry inhibitor; FNTA, farnesyltransferase, CAAX box, subunit alpha; FNTB, farnesyltransferase, CAAX box, subunit beta; HTAs, host-targeting agents; LN, lonafarnib; Mig.cDCs, migratory cDCs; NK cells, natural killer cells; pDCs, plasmacytoid DCs; scRNA-seq, single-cell RNA sequencing; SLC10A1, solute carrier family 10 member 1; SLGN, selgantolimod; TLR8, Toll-like receptor 8; UMAP, uniform manifold approximation and projection.

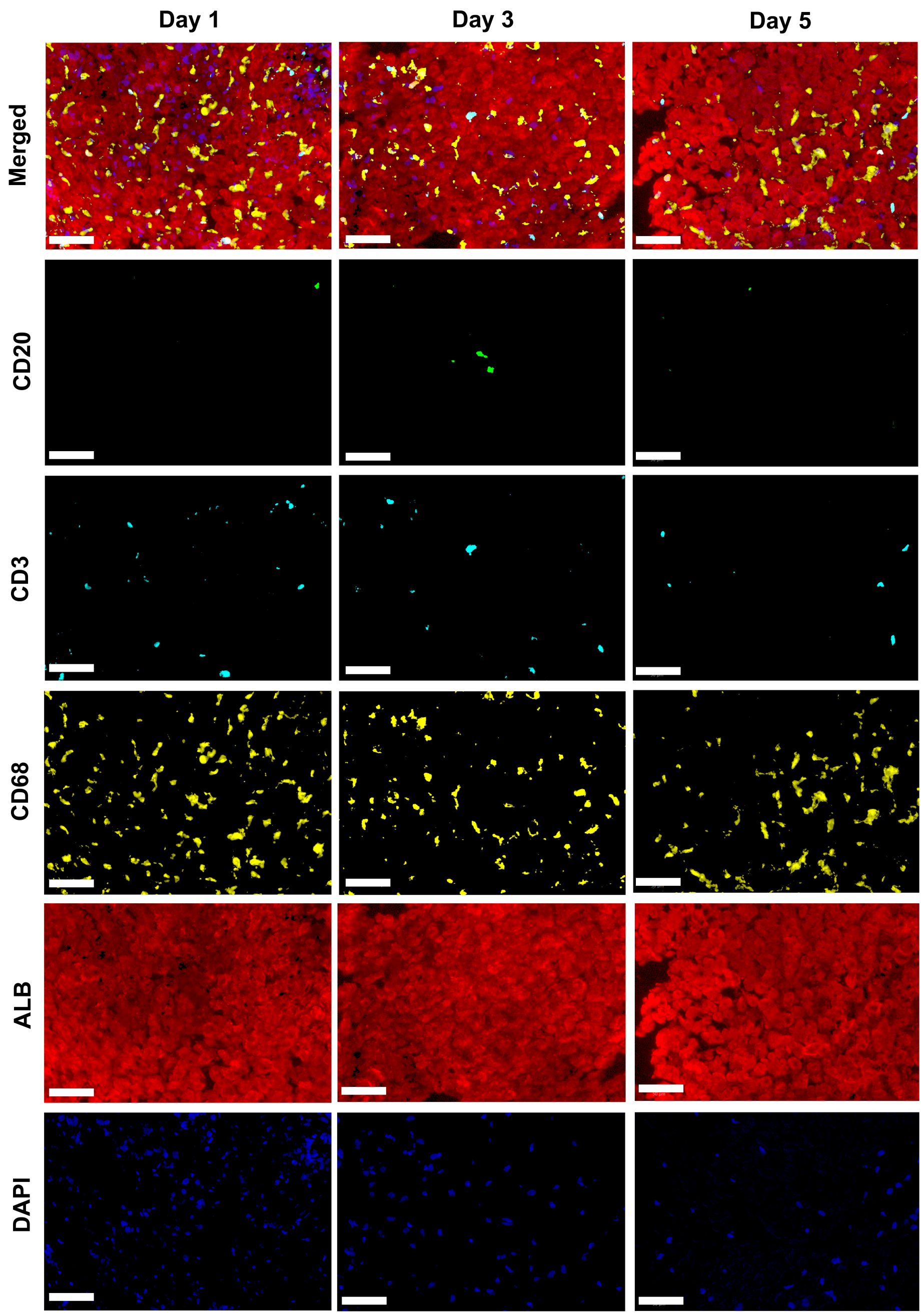

**Fig. S4. Parenchymal and immune populations present in PCLS.** Immunofluorescence image of human PCLS cultured for 1, 3, and 5 days post-cutting and stained for CD20 (B cells), CD3 (T cells), CD68 (general myeloid), ALB (hepatocytes), and DAPI (nuclear DNA). Scale bar represents 50 µm. ALB, albumin; PCLS, precision-cut liver slices.

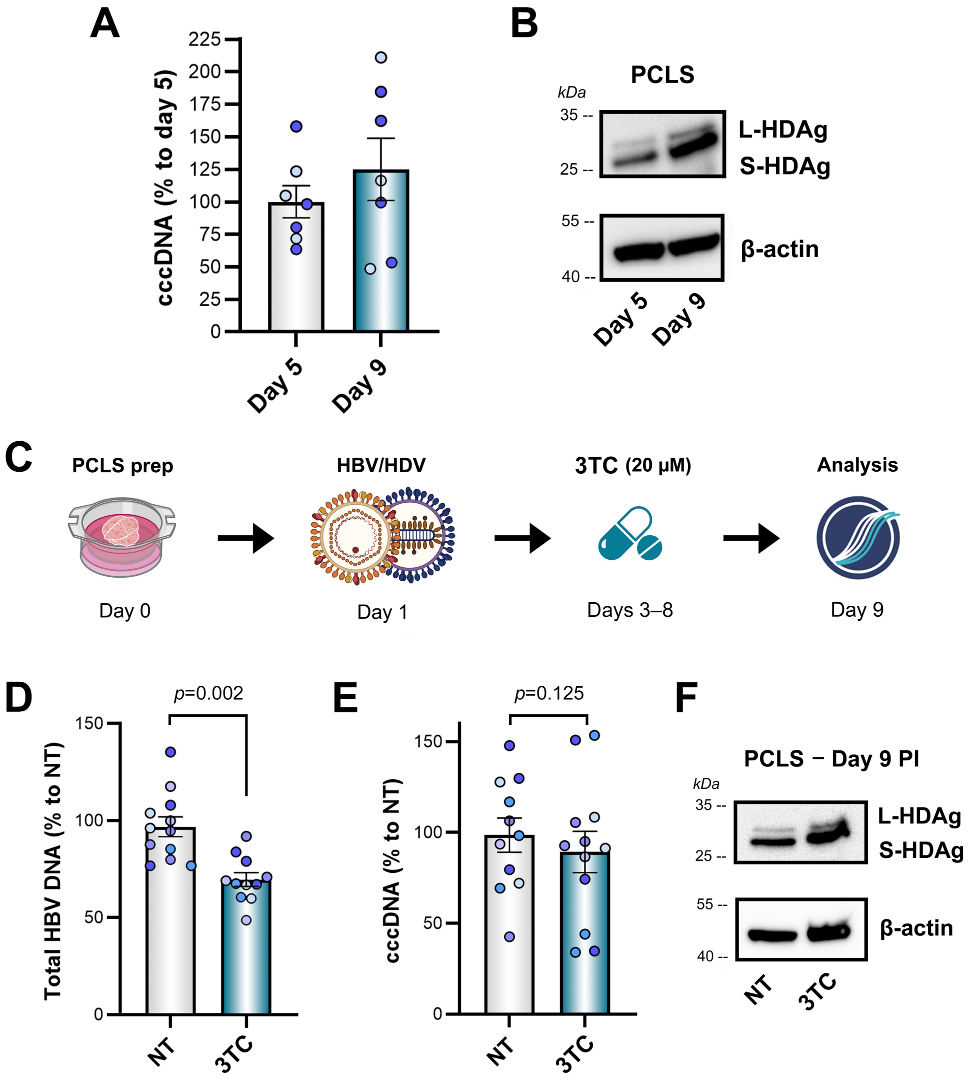

**Fig. S5. Treatment with 3TC decreases total HBV DNA in the PCLS model.** (**A**) Quantification of HBV cccDNA by qPCR in HBV/HDV co-infected PCLS at days 5 and 9 post-infection ($n = 2$). (**B**) Detection by western blot of intracellular large and small HDAg in HBV/HDV-infected PCLS at days 5 and 9 post-infection. (**C**) Experimental protocol consisting of the preparation of PCLS, their co-infection with HBV/HDV for a nine-day period, and the use of 3TC (lamivudine, 20 µM) post-infection in order to evaluate its antiviral effect by the quantification of viral parameters. Image created in BioRender. (**D–E**) Quantification of (D) total HBV DNA and (E) cccDNA by qPCR in HBV/HDV co-infected PCLS ($n = 5$, Mann-Whitney test). Bars represent mean ± SEM. Dot colors indicate individual liver tissue donors ($n$). (**F**) Detection by western blot of intracellular large and small HDAg in HBV/HDV-infected PCLS. cccDNA, covalently closed circular DNA; HBV, hepatitis B virus; HDAg, hepatitis delta antigen; HDV, hepatitis delta virus; PCLS, precision-cut liver slices.

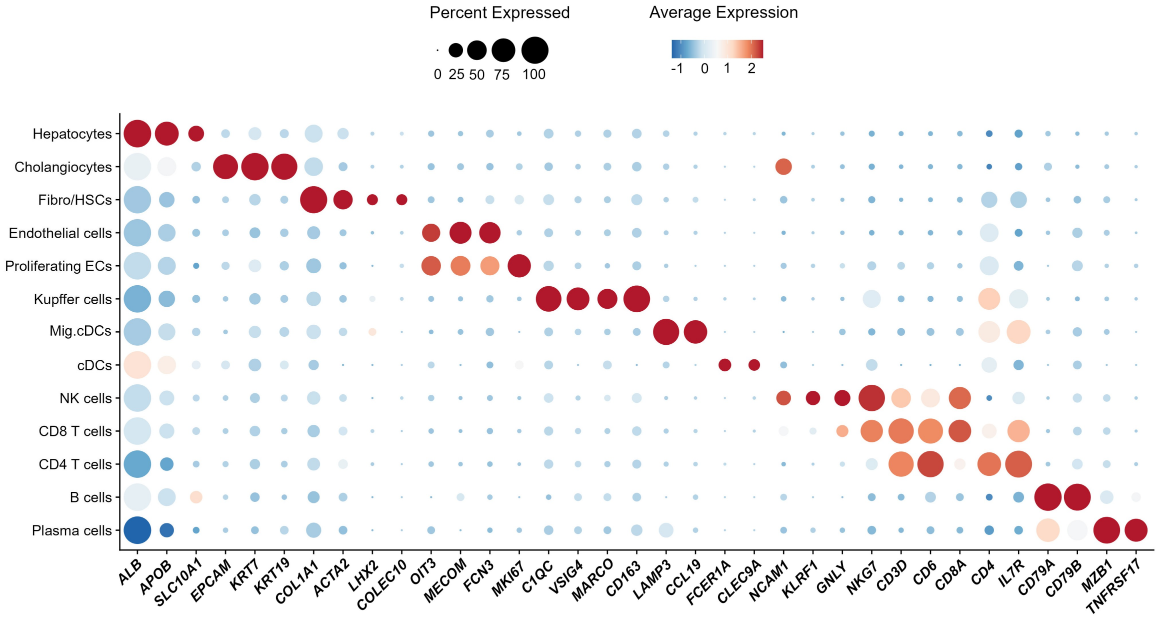

**Fig. S6. Identification of hepatic cell populations present in human PCLS.** Dotplot showing the expression levels of canonical cell type markers employed for the annotation of each cell type, as quantified by scRNA-seq. ACTA2, actin alpha 2, smooth muscle; ALB, albumin; APOB, apolipoprotein B; C1QC, complement C1q C chain; CCL19, C-C motif chemokine ligand 19; cDCs, classical dendritic cells; CLEC9A, C-type lectin domain containing 9A; COL1A1, collagen type I alpha 1 chain; COLEC10, collectin subfamily member 10; ECs, endothelial cells; EPCAM, epithelial cell adhesion molecule; FCER1A, Fc epsilon receptor Ia; FCN3, ficolin 3; GNLY, granulysin; HSCs, hepatic stellate cells; IL7R, interleukin 7 receptor; KLRF1, killer cell lectin like receptor F1; KRT, keratin; LAMP3, lysosome associated membrane protein 3; LHX2, LIM homeobox 2; MARCO, macrophage receptor with collagenous structure; MECOM, MDS1 and EVI1 complex locus; Mig.cDCs, migratory cDCs; MKI67, marker of proliferation Ki-67; MZB1, marginal zone B and B1 cell specific protein; NCAM1, neural cell adhesion molecule 1; NK cells, natural killer cells; NKG7, natural killer cell granule protein 7; OIT3, oncoprotein induced transcript 3; PCLS, precision-cut liver slices; scRNA-seq, single-cell RNA sequencing; SLC10A1, solute carrier family 10 member 1; TNFRSF17, TNF receptor superfamily member 17; VSIG4, v-set and immunoglobulin domain containing 4.

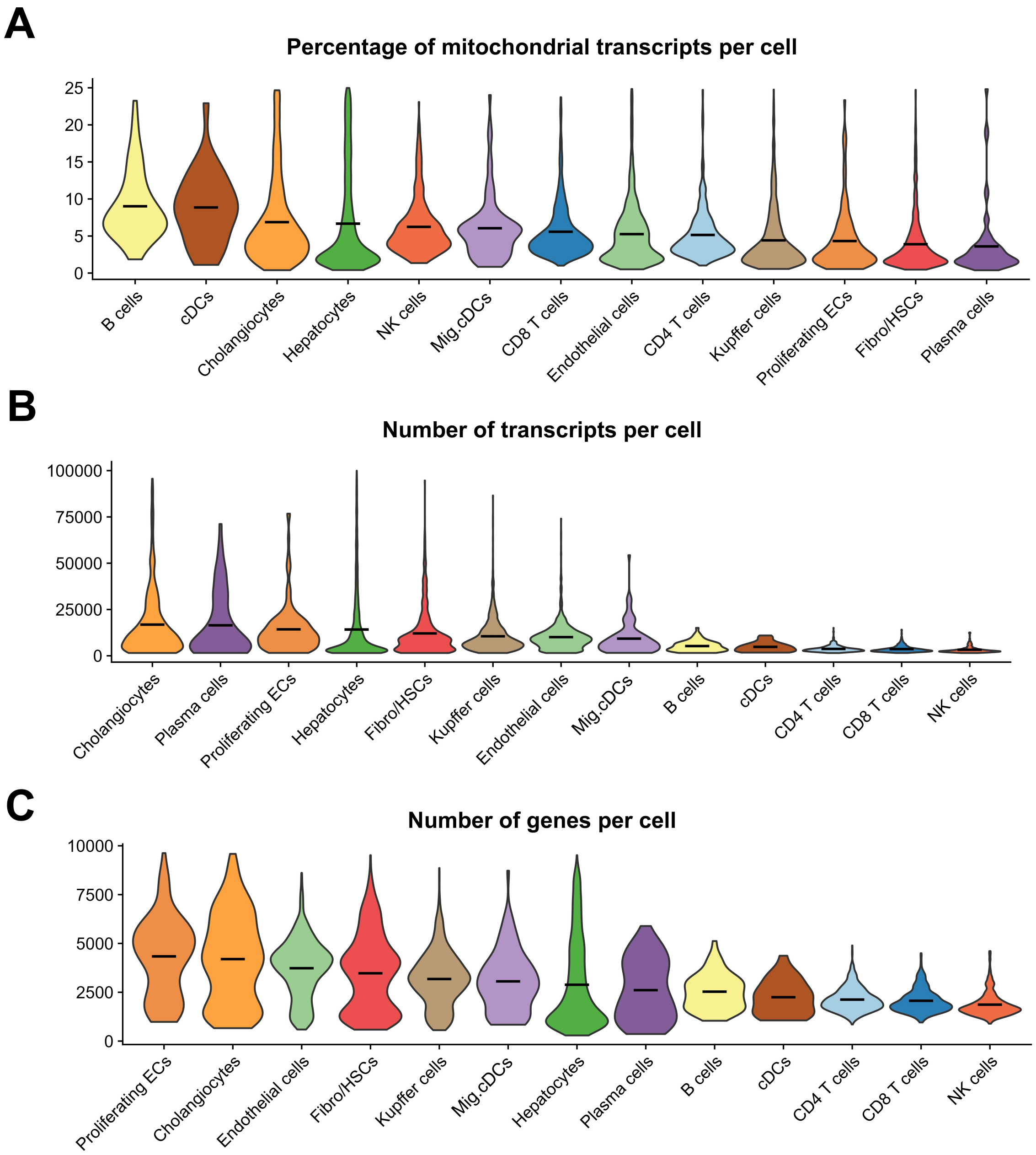

**Fig. S7. QC parameters employed to select high-quality cells. (A–C)** Violin plots depicting (A) the percentage of mitochondrial transcripts/cell (<25%), (B) total number of transcripts/cell (>1500, <99999), and (C) number of genes/cell (>200), as quantified by scRNA-seq. Violin plots represent mean values. cDCs, classical dendritic cells; ECs, endothelial cells; HSCs, hepatic stellate cells; Mig.cDCs, migratory cDCs; NK cells, natural killer cells; PCLS, precision-cut liver slices; scRNA-seq, single-cell RNA sequencing.

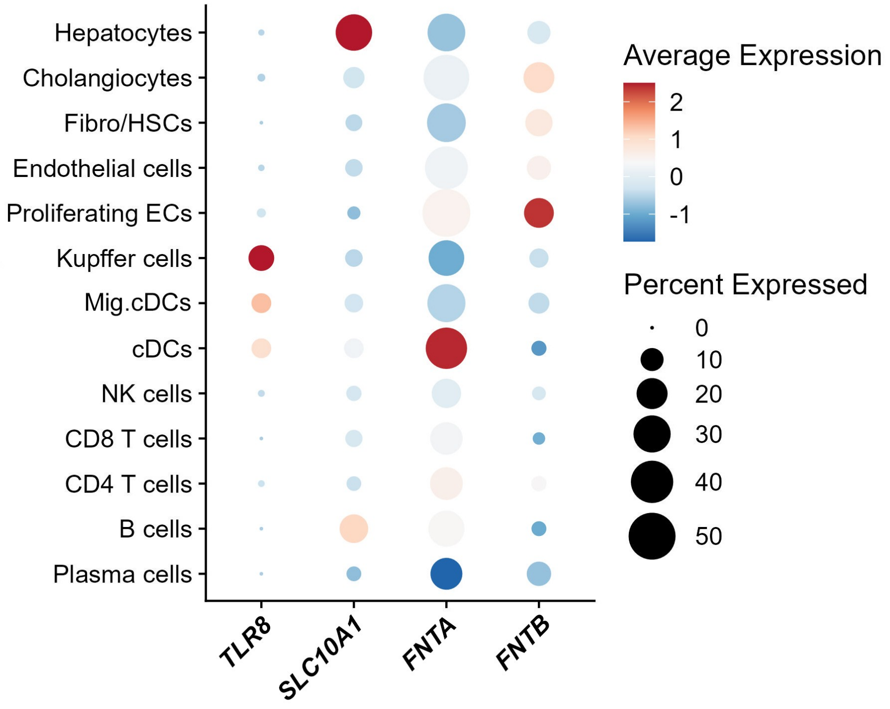

**Fig. S8. Expression of the cellular targets related to the HTAs employed for PCLS treatment.** Cell populations in PCLS showing expression of *SLC10A1* (NTCP, EI target), *TLR8* (SLGN target), *FNTA* and *FNTB* (LN targets), as quantified by scRNA-seq. cDCs, classical dendritic cells; ECs, endothelial cells; EI, entry inhibitor; FNTA, farnesyltransferase, CAAX box, subunit alpha; FNTB, farnesyltransferase, CAAX box, subunit beta; HSCs, hepatic stellate cells; HTAs, host-targeting agents; LN, lonafarnib; Mig.cDCs, migratory cDCs; NK cells, natural killer cells; PCLS, precision-cut liver slices; scRNA-seq, single-cell RNA sequencing; SLC10A1, solute carrier family 10 member 1; SLGN, selgantolimod; TLR8, Toll-like receptor 8.

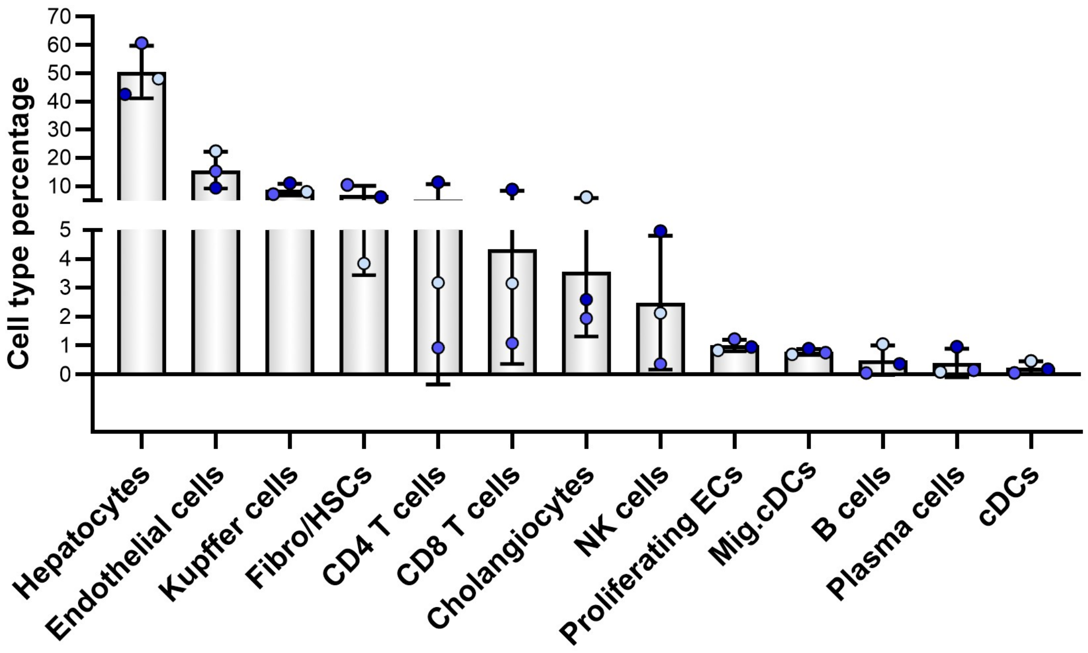

**Fig. S9. Cell population proportions present within PCLS for each individual tissue donor.** Percentage of individual cell populations identified by scRNA-seq for each tissue donor ($n$ = 3). Bars represent mean ± SD. Dot colors indicate individual liver tissue donors ($n$). cDCs, classical dendritic cells; ECs, endothelial cells; HSCs, hepatic stellate cells; Mig.cDCs, migratory cDCs; NK cells, natural killer cells; PCLS, precision-cut liver slices; scRNA-seq, single-cell RNA sequencing.

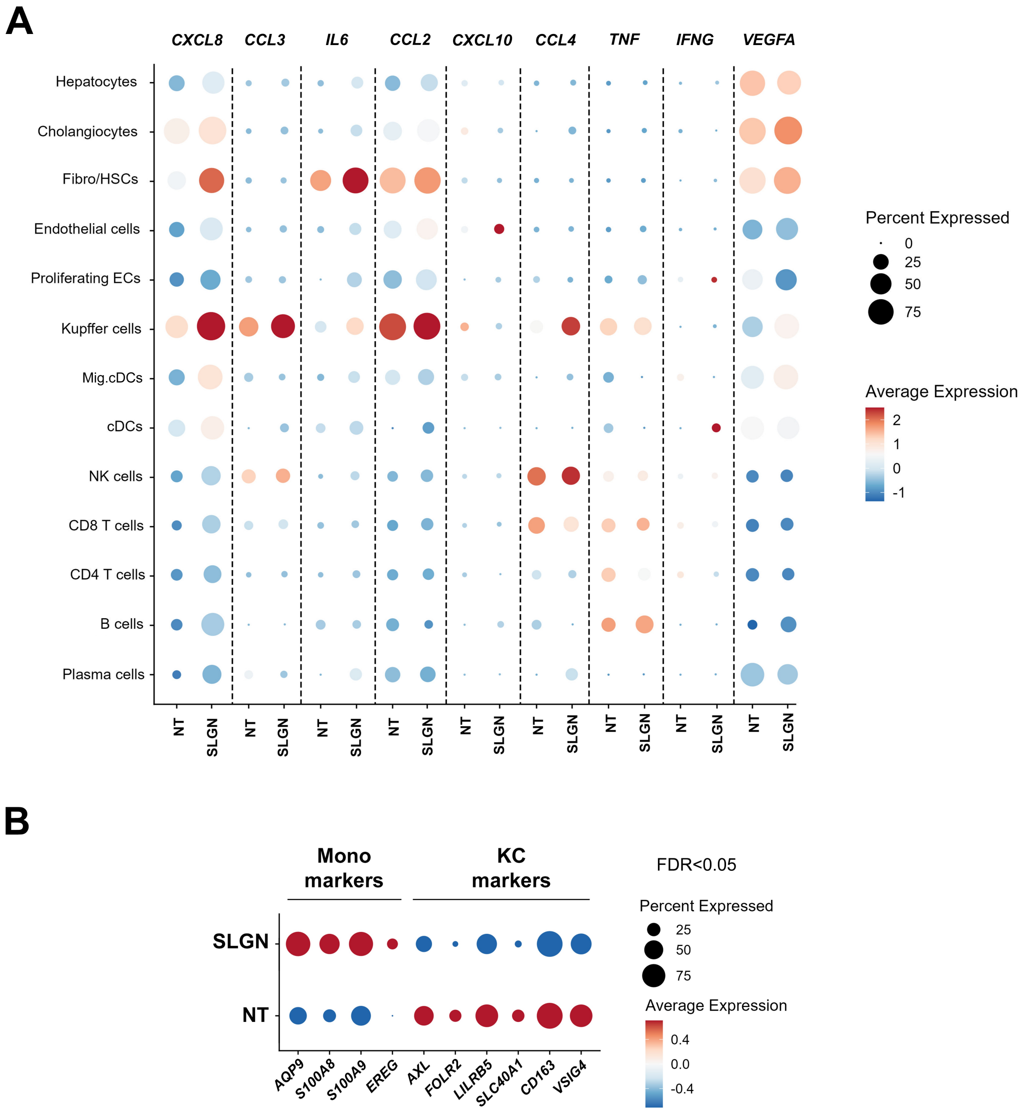

**Fig. S10. Expression of inflammatory cytokines and KC differentiation markers in response to SLGN. (A)** Expression of inflammatory cytokines at the mRNA level from each cell type within PCLS following SLGN treatment (*n* = 3, 1 μM, 48 h). **(B)** Expression of differentiation markers at the mRNA level from KCs within PCLS following SLGN treatment, as quantified by scRNA-seq. AQP9, aquaporin 9; AXL, AXL receptor tyrosine kinase; CCL, C-C motif chemokine ligand; cDCs, classical dendritic cells; CXCL, C-X-C motif chemokine ligand; ECs, endothelial cells; EREG, epiregulin; FOLR2, folate receptor beta; HSCs, hepatic stellate cells; IFNG, interferon gamma; KCs, Kupffer cells; LILRB5, leukocyte immunoglobulin like receptor B5; Mig.cDCs, migratory cDCs; NK cells, natural killer cells; PCLS, precision-cut liver slices; S100A, S100 calcium binding protein A; scRNA-seq, single-cell RNA sequencing; SLC40A1, solute carrier family 40 member 1; SLGN, selgantolimod; TNF, tumor necrosis factor; VEGFA, vascular endothelial growth factor A; VSIG4, v-set and immunoglobulin domain containing 4.

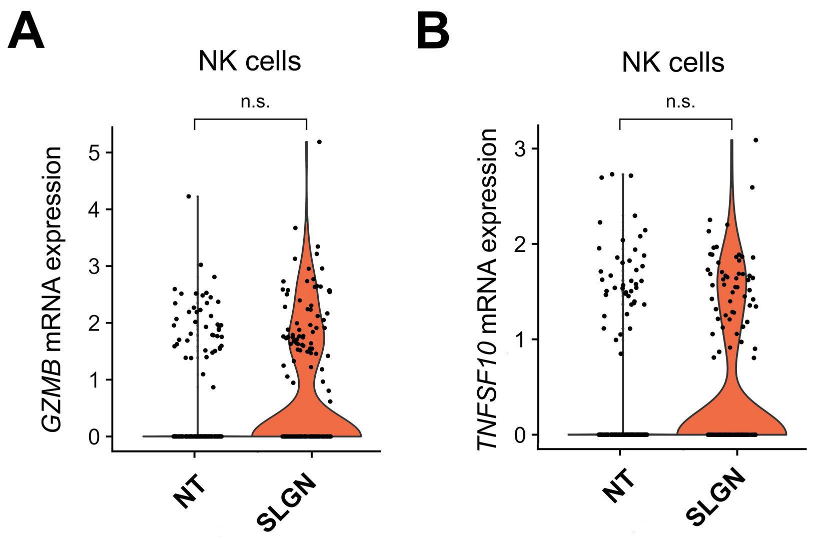

**Fig. S11. Increase of NK activation markers in response to SLGN. (A–B)** Expression levels of (A) *GZMB* and (B) *TNFSF10* (TRAIL) in NK cells from PCLS treated with SLGN (*n* = 3, 1 µM, 48 h). GZMB, granzyme B; NK cells, natural killer cells; PCLS, precision-cut liver slices; SLGN, selgantolimod; TNFSF10, tumor necrosis factor (ligand) superfamily 10.

| Donor | Age | Sex | BMI | Fib/Cir | Steatosis | Diabetes | HBV | HCV | Alcoholism | Diagnosis |
|---|---|---|---|---|---|---|---|---|---|---|
| 1* | 73 | Male | 30.10 | No | No | No | No | No | No | Choroidal melanoma |
| 2* | 39 | Male | 21.04 | No | No | No | No | No | No | Colorectal cancer |
| 3* | 74 | Female | 32.05 | No | No | Yes | No | No | No | Colorectal cancer |
| 4 | 72 | Male | 28.67 | No | No | No | No | No | No | Colorectal cancer |
| 5 | 68 | Female | 17.85 | No | No | Yes | No | No | No | Ciliochoroidal melanoma |
| 6 | 71 | Male | 26.89 | No | No | No | No | No | No | Colorectal cancer |
| 7 | 43 | Male | 25.76 | No | No | No | No | No | No | Choroidal melanoma |
| 8 | 73 | Female | 19.40 | No | No | No | No | No | No | Colorectal cancer |
| 9 | 60 | Female | 28.90 | No | No | Yes | No | No | No | Colorectal cancer |
| 10 | 61 | Male | 23.00 | No | No | No | No | No | No | Colorectal cancer |
| 11 | 62 | Male | 19.00 | No | No | No | No | No | Yes | Colorectal cancer |
| 12 | 49 | Male | 26.00 | No | No | No | No | No | No | Colorectal cancer |
| 13 | 57 | Male | 34.00 | No | No | No | No | No | Yes | Colorectal cancer |

**Table S1. Clinical data from liver tissue donors employed for the preparation of PCLS.** BMI, body mass index; Cir, cirrhosis; Fib, fibrosis; HBV, hepatitis B virus; HCV, hepatitis C virus. *PCLS from liver resection donors characterized by scRNA-seq.

| Primer name: | Sequence (5' -> 3'): |
|---|---|
| cccDNA-FW | CCGTGTGCACTTCGCTTCA |
| cccDNA-BW | GCACAGCTTGGAGGCTTGA |
| cccDNA probe | (6FAM)CATGGAGACCACCGTGAACGCCC(BBQ) |
| Total HBV RNA/DNA | Vi03453406_s1 (Thermo Fisher Scientific) |
| Total HDV RNA-FW | CGGGCCGGCTACTCTTCT |
| Total HDV RNA-BW | AAGGAAGGCCCTCGAGAACA |
| HDV-genomic-FW | Biotin-GCGCCGGCYGGGCAAC |
| HDV-anti-genomic-BW | Biotin-TTCCTCTTCGGGTCGGCATG |
| *HBB* | Hs00758889_s1 (Thermo Fisher Scientific) |
| *GUSB* | Hs99999908_m1 (Thermo Fisher Scientific) |

**Table S2. Primer sequences.** Biotinylated primer sequences for the detection of genomic and anti-genomic HDV RNA were obtained from Giersch *et al.*, Sci Rep. 2017.[2] cccDNA, covalently closed circular DNA; GUSB, glucuronidase beta; HBB, hemoglobin subunit beta; HBV, hepatitis B virus; HDV, hepatitis delta virus.
